# Modeling Cell Shape Diversity Arising From Complex Rho Gtpase Dynamics

**DOI:** 10.1101/561373

**Authors:** Cole Zmurchok, William R. Holmes

## Abstract

It is well known that cells exhibit a variety of morphologically distinct responses to their environments that manifest in their cell shape. Some protrude uniformly to increase substrate contacts, others are broadly contractile, some polarize to facilitate migration, and yet others exhibit mixtures of these responses. Prior imaging studies have identified a discrete collection of shapes that the majority of cells display and have demonstrated links between those shapes and activity levels of the cytoskeletal regulators Rho GTPases. Here we use a novel computational modeling approach to demonstrate that well known Rho GTPase signaling dynamics naturally give rise to this diverse but discrete (rather than continuum) set of morphologies. Specifically, the combination of auto-activation and mutually-antagonistic crosstalk between GTPases along with the conservative membrane (un)binding dynamics readily explain at least 6 of the 7 commonly observed morphologies. We further use this methodology to map the entire parameter space of this model and show that in appropriate regimes, individual parameter sets give rise to a variety of different morphologies. This provides an explanation for how seemingly similar cells of the same fate derived from the same population can exhibit a diverse array of cell shapes in imaging studies. These results thus demonstrate that Rho GTPases form the core of a cytoskeletal regulatory system governing cell shape, further supporting the picture that they act as a central signaling hub determining how cells respond to their environmental context.

## 1. INTRODUCTION

It is well documented that similar cells from a single population can exhibit different morphologies (shape in particular [1,2]) and behaviors (amoeboid versus mesenchymal migration [3]). Where does this diversity come from? One possibility is that the cells are functionally different, possibly differing in their expression of proteins or cytoskeletal factors that determine the cells’ structure. Another is that the cells really are similar but that we are observing their exploration of a complex morphological state space. In this article we utilize a new computational modeling approach to assess whether and to what extent the dynamics of a crucial class of cytoskeletal regulators, the Rho GTPases, may explain this diversity.

Despite using the same (or similar) molecular machinery to regulate and remodel the cytoskeleton, cells exhibit a variety of different morphologies. For example, cultured *Drosophila* haemocytes can be classified as a heterogeneous mixture of five discrete shapes (normal, elongated, large-ruffled, teardrop, large-smooth) [4]. *Drosophila* BG-2 cells were shown to exhibit seven “shapes” varying in size and type of polarization [2]. Melanoma cells in 3D matrices can exhibit either mesenchymal or amoeboid modes of migration and also give demonstrate a mixture of six different shapes (star, spindle, teardrop, ellipse, small round, large round) [5]. Given the recurrence of a relatively restricted set of morphologies across cell types, it has been proposed [6] that there may be a common mechanism underpinning cell shape determination that gives rise to a relatively simple morphological landscape and that genetic and environmental forces tune this landscape. Here we demonstrate that Rho GTPase dynamics alone has the potential to generate an expansive set of shape dynamics and that these proteins’ dynamics may form the foundation of such a common mechanism.

Numerous cell characteristics such as shape, protrusiveness, polarization [2], and mode of migration [3,7] have been causally linked to Rac1 and RhoA GTPase activity levels (henceforth Rac and Rho). These proteins are known to be central regulators of cytoskeletal remodeling processes that determine cells’ morphologies. Broadly, Rac activity is associated with actin polymerization and cellular protrusion while Rho activity is associated with actomyosin contraction [8,9]. For this reason, GTPase activity levels have been linked to cell shape (Figure 1(a)). Cells that have high levels of Rac and little Rho are spread out and flat while those that have low levels of Rac and high Rho are contracted [2]. Cell polarity and migration is, in many cases, characterized by spatially distinct zones of cell behavior and GTPase signaling: a Rac-dominated protrusive front and Rho-dominated contractile rear [10, 11, 12, 13] (reviewed in [14]).

**FIGURE 1.**
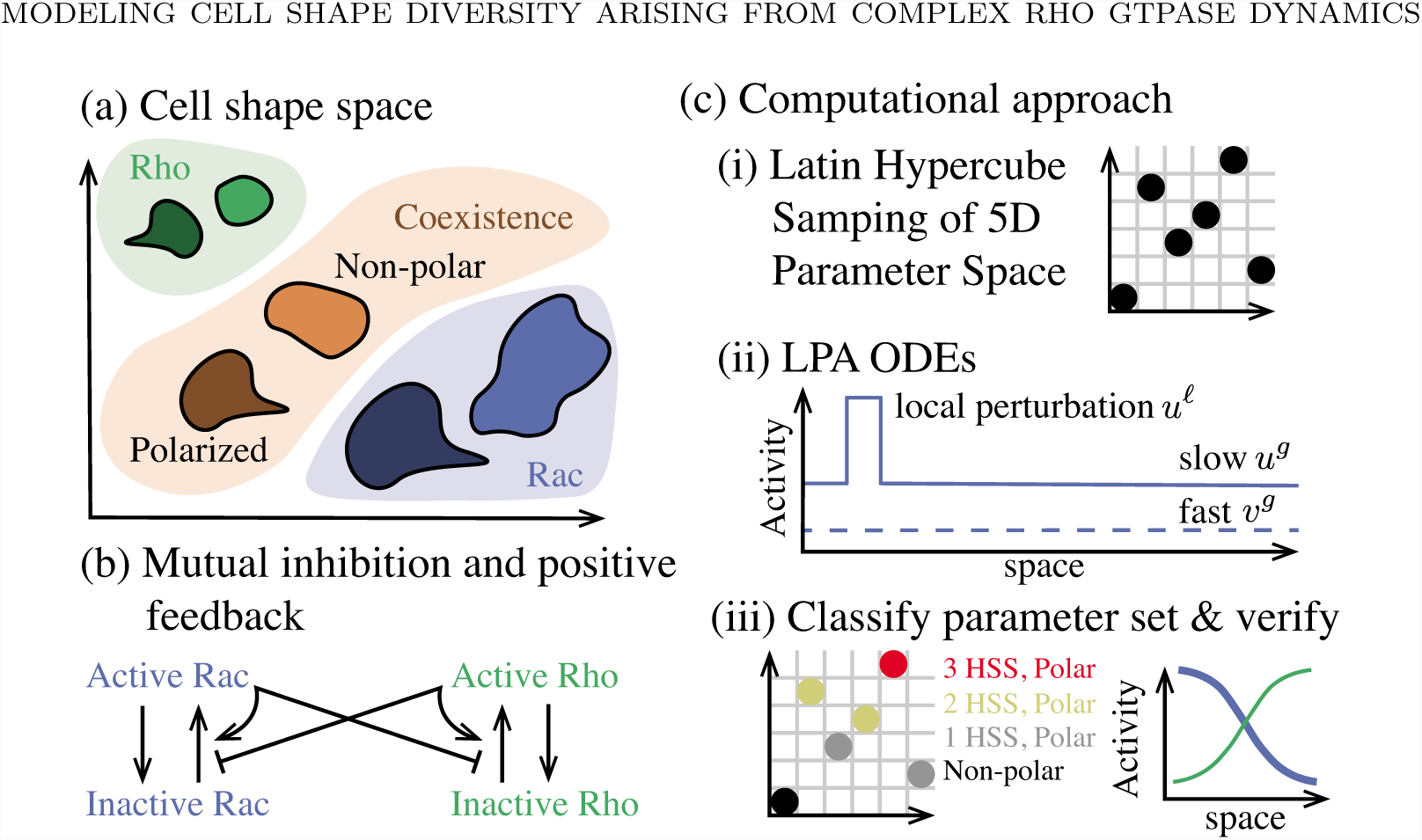
Overview of the biological context, model, and computational approach. **(a)** Schematic of cell morphologies and their link to Rac and Rho regulation [2]. Generally Rac dominated cells tend to be protrusive and spread out while Rho dominated cells tend to be contractile and smaller. **(b)** Schematic of mutual-antagonism, auto-activation, and conservation model investigated here. Active (respectively inactive) proteins are assumed to be bound (resp. unbound) from the cell membrane and thus slow (resp. fast) diffusing. **(c)** Overview of computational approach to assess this models dynamics and parameter space structure. (i) Latin Hypercube Sampling is used to obtain a computationally efficient sample of a five-dimensional (5D) parameter space. (ii) The Local Perturbation Analysis (LPA) is used to approximate the partial differential equation (PDE) system by a system of ordinary differential equations (ODEs) that describe the evolution of a local perturbation *u*^*ℓ*^ against background slowly diffusing *u*^*g*^ and quickly diffusing *v*^*g*^ variables.(iii) Well-mixed analysis and LPA are used to characterize the properties of each parameter set (capacity for multistability and polarization) and PDE simulations are used for confirmation and further analysis.

Due to their central role in regulating the cytoskeleton, these proteins have been the source of intense investigation since their discovery. This has led to a picture where Rho GTPases sit at the center of larger signaling network involving interactions with ligand gradients [9], the extracellular matrix [15], and even mechanical forces [16] and membrane tension [17]. In addition to interacting with these external signaling systems, Rho GTPases have been demonstrated to mutually antagonize each other [18,19] as well as auto-activate themselves [20]. Here we use mathematical modeling to explore the consequences of these Rho GTPase feedbacks (independent of the larger signaling web) on the types of spatial distributions of protein activation that form the foundation of cell shape determination.

Mathematical and computational modeling have long been used to investigate the role of GTPases on cell dynamics (see for example [21, 22, 23, 24, 25, 26, 27, 28] along with [29, 30, 31, 32] for reviews). Recent efforts over the last decade have led to the hypothesis that two critical aspects of GTPase dynamics are responsible for their ability to generate the kinds of spatial patterns of activation that are a prerequisite for polarity. The “wave pinning” model [33, 34, 35] of GTPase dynamics demonstrated that the combination of nonlinear feedback (such as auto-activation or mutual-antagonism) along with mass conservation yields stable, polarized spatial distributions of protein activation. A recent extension [36] has shown that the combination of non-linearity and mass conservation can also produce protein distributions indicative of different morphologies. This work however considered a somewhat restricted model of GTPase dynamics and only explained a subset of observed morphologies. Here we extend this work and use computational modeling to demonstrate that known GTPase dynamics explain a wider diversity of cell shapes than previously shown.

In this article, we study a spatial model of Rho GTPase dynamics encoding the auto-activation, mutual-antagonism, and mass conservation dynamics that have been observed in numerous prior experimental studies. A central challenge in studying the spatial dynamics of complex, nonlinear models such as this is the difficulty in systematically studying the partial differential equations (PDEs) that mathematically describe those dynamics. We thus develop and validate a new computational approach based on the Local Perturbation Analysis (LPA) [37, 38, 39] to efficiently and systematically study the dynamics of this model in the whole parameter space. Using this approach, we find that the combination of mutual-antagonism, auto-activation, and conservation leads to a variety of spatial GTPase activity profiles that give rise to at least six distinct previously observed cell morphologies (illustrated in Figure 1(a)). Furthermore, we show that this system can generate a wide variety of spatial protein activation profiles (linked to different morphologies) from a single state (e.g. parameter set). Thus much of the variance in cell shape within a population of cells could be a result of those cells exploring the morphological landscape, rather than being a reflection of intrinsic differences between the cells (e.g. gene expressions). In conclusion, this work demonstrates that Rho GTPase dynamics may form the foundation of the mechanism responsible for cell shape determination in a wide variety of cells.

## 2. RESULTS

Our goal here is to study the link between Rho GTPase (Rac and Rho specifically) dynamics and the diverse cell shapes observed in cellular populations (illustrated in Figure 1(a)). Towards this end, we study a spatiotemporal model of GTPase dynamics that incorporates the primary known characteristics of Rac and Rho dynamics: auto-activation, mutual-antagonism, and conservation (Figure 1(b)). Numerous past investigations have studied the temporal dynamics of this canonical mutual-antagonism and auto-activation system [40, 41, 42]. This study differs from such prior studies of this motif in two ways. First, it focuses on spatial dynamics in the context of cell polarity. Second, we incorporate biochemical conservation, which is known to be a vital aspect of Rho GTPase dynamics that leads to the formation of new spatiotemporal behaviors [33, 34, 35, 43]. In order to characterize the spatiotemporal behaviors of this mutual-antagonism, auto-activation, and conservation system in a systematic fashion, we use a multi-faceted approach combining well-mixed analysis, Local Perturbation Analysis (LPA) [31, 37, 38, 39, 44], and PDE simulation with unsupervised parameter space screening (Figure 1(c)).

### 2.1. Model background and description

This study is inspired by, and bridges, two primary bodies of literature regarding cell polarity regulation and multistability in gene regulation. Studies of cell polarity [33, 34, 35, 36] have revealed that the combination GTPase conservation and either mutual-antagonism or auto-activation leads to a novel type of dynamic referred to as “wave pinning.” Figure 2(a) illustrates (using a relatively new method, the LPA, which is described further in the §4) that the combination of mutual-antagonism and GTPase conservation can lead to a diverse array of spatial Rho GTPase activation profiles including uniformly Rac activated, uniformly Rho activated, or polarized states. This literature has however not considered the consequences of both auto-activation and mutual-antagonism jointly acting.

**FIGURE 2.**
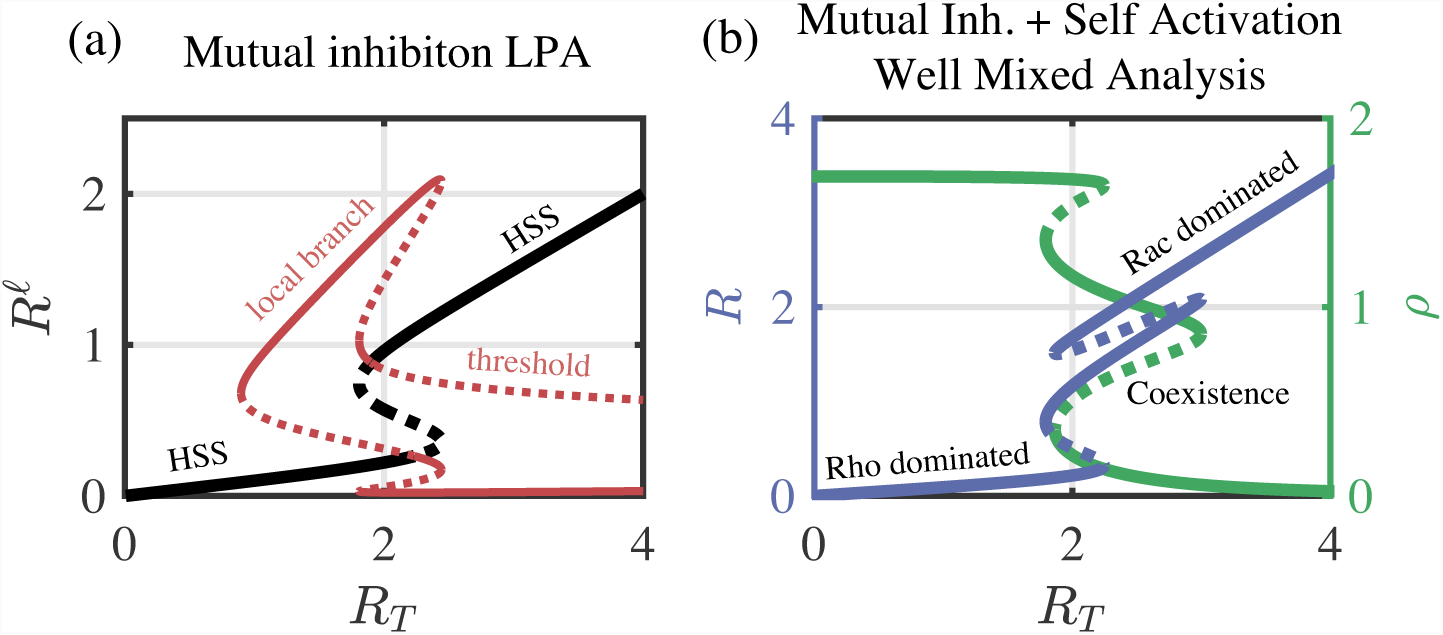
Bistability, tristability and polarization. **(a)** LPA bifurcation analysis of the mutual-antagonism model (*a* = 0, similar to results in [36]). The global (black curve) and local solution branches (red curve) are shown along with their stability (solid, stable; dashed, unstable). A region of well-mixed bistability is enclosed in a larger region where stimulus induced polarization is possible (via a perturbation across a threshold). Parameters are *b* = 1, *c* = 0, *s* = 0.5, *n* = 3, and *ρ*_*T*_ = 2. **(b)** Well-mixed bifurcation analysis of the mutual-antagonism and auto-activation model (*a* > 0). Tristability is possible with both auto-activation and mutual-antagonism. The middle branch is referred to as the coexistence HSS, where both Rac and Rho activity are at moderate levels. Parameters are *a* = 1.8, *b* = 4, *s* = 0.5, *n* = 3, and *ρ*_*T*_ = 2.

On the other hand, the simple *tristable* circuit comprised of mutual-antagonism and auto-activation has been extensively studied in gene expression literature [40, 41, 42]. Figure 2(b) illustrates a typical well-mixed bifurcation diagram showing the well known tristability this system gives rise to. Much of this literature has however focused on temporal aspects of this system and, since much of this work is related to gene regulation, does not consider the consequences of biochemical conservation.

We study the spatiotemporal consequences of all three of these elements (mutual-antagonism, auto-activation, and biochemical conservation) regulating cell dynamics, all of which are pertinent to GTPase dynamics. Toward this end, we thus consider a PDE model of this system utilizing standard Hill function representations of feedbacks:

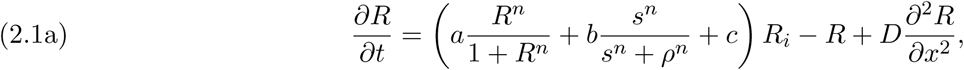

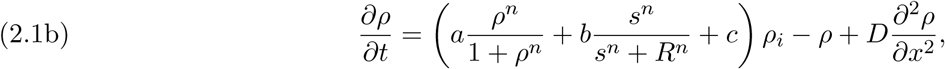

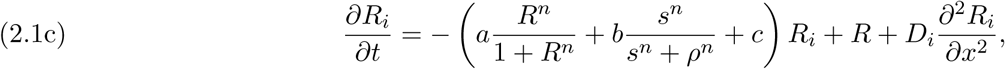

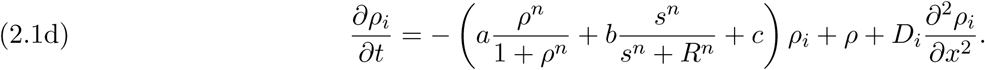

Here, we model the GTPase activity in a one dimensional domain with no-flux boundary conditions, representing an averaged cross section of a cell. *R*(*x, t*) and *ρ*(*x, t*) are the slowly diffusing (*D*) membrane-bound active forms of Rac and Rho GTPase, while *R*_*i*_(*x, t*) and *ρ*_*i*_(*x, t*) are the freely diffusing (*D*_*i*_) cytosolic inactive forms. Since the GTPases simply switch between active and inactive forms with no production or degradation on the time scales of interest, the total amount of each GTPase is conserved: 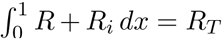 and 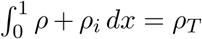. The activation rates (terms in parenthesis) capture the feedbacks between GTPases: *a* is the strength of the auto-activation, *b* is the strength of the mutual-antagonism switch, and *c* is the baseline activation rate of GTPase. For simplicity, the parameters *a, b, c*, and *s* are assumed to be the same for Rac and Rho and the system has been non-dimensionalized so that the half maximum of the auto-activation Hill function is 1 and the deactivation rates are 1. See §4 for full model details.

### 2.2. Computational approach

The goal of this study is to broadly characterize the different steady-state spatial distributions of activated GTPase concentrations arising from this model. Rather than simply choose a representative parameter set to study, we have developed an efficient approach to map spatiotemporal regimes of behavior of this system in parameter space. This is a multi-faceted approach that combines the use of well-mixed analysis, a relatively new non-linear perturbation analysis method (the Local Perturbation Analysis or LPA), and PDE simulation to perform an unsupervised screen of parameter space.

To characterize the model behavior, we use the Latin Hypercube sampling method to generate a computationally efficient sample of high-dimensional parameter space (Figure 1(c)(i)). For each given parameter set of this model that is considered, we utilize three methods of analysis that provide increasingly more refined information about the model’s dynamics. (1) Well-mixed analysis is used to determine what kinds of spatially homogeneous states are present. For example, a uniformly Rac (or Rho) dominated state would be associated with a spread (or contracted) morphology. (2) The Local Perturbation Analysis is used to rapidly assess whether non-uniform steady-states (e.g. polarization) are expected. This is a highly efficient approximate non-linear perturbation analysis that predicts whether the model will respond to a sufficiently large perturbation (Figure 1(c)(ii)) and give rise to a spatial pattern. (3) Finally, PDE simulation will be used to check for and determine the nature of pattern formation. This multi-faceted approach will combine all three methods into a coherent computational framework. For a detailed description of each of these methods, see §4.

### 2.3. The combination of mutual-antagonism, auto-activation, and biochemical conservation leads to a wide array of spatial GTPase activation profiles that match known cell morphologies

Before broadly characterizing the parameter space dynamics of this model, we first illustrate the types of states this system elicits and discuss how those map onto previously observed cell morphologies. It is well known that, for a single model parameter set, a system comprised of mutual-antagonism and auto-activation can elicit three or even four distinct, stable well-mixed states. Alternatively, either of these feedbacks in conjunction with conservation is known yield coexistence of uniform and polarized steady-states [36]. We thus hypothesize that the combination of all three may yield a variety of diverse stable spatial protein distributions, each of which correspond to a distinct cell morphology.

PDE simulation results for select parameter sets show that this is indeed the case (Figure 3). Panels (a)-(d) demonstrate that for a single parameter set, three possible well mixed states along with a polarized state are possible, all of which are stable. Each of these steady-states would be linked to a different morphology (e.g. morphologies cell populations from [1, 2, 5]), indicated by the inset cartoons. Given the propensity for Rac to generate protrusion and Rho contraction, Rac and Rho dominated states respectively would be associated with large, spread cells and small, contracted cells. Alternatively, the state where both are at moderate levels would be associated with a kind of balanced rest state that is neither overly protrusive or contractile. Finally, the heterogeneous state (c) would be associated with a polarized cell that protrudes on one side and contracts on the other. Importantly, this simple system can give rise to each of these protein distributions for a single parameter set.

**FIGURE 3.**
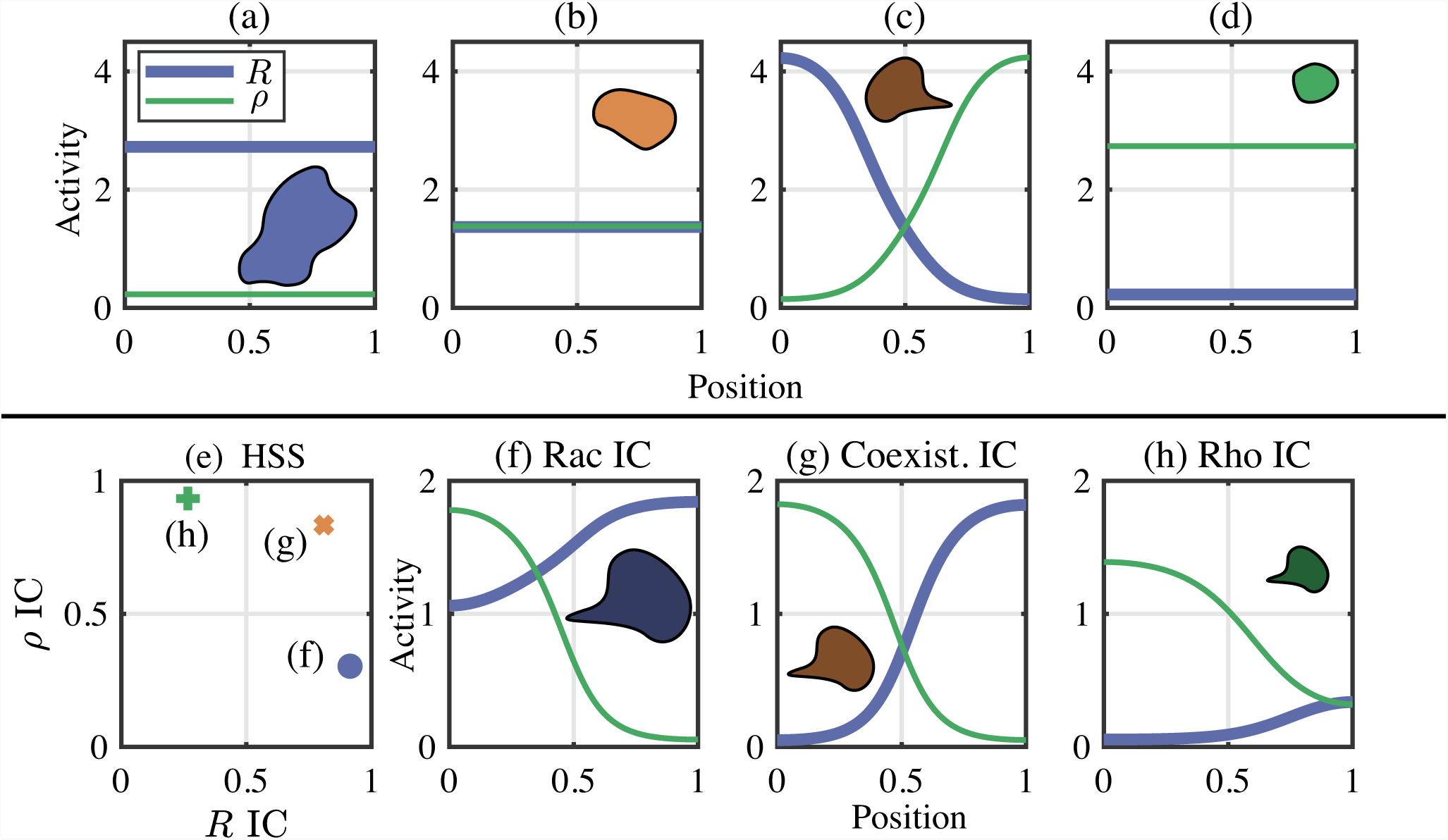
Example PDE simulations of diverse protein activation states. **(a)** Rac dominated HSS. **(b)** Coexistence HSS. **(c)** Polarized cell obtained from a local stimulus applied to the coexistence HSS. **(d)** Rho dominated HSS. Panels (a)-(d) are all simulated from the same parameter set: *a* = 0.4898, *b* = 1.2034, *c* = 0.0415, *R*_*T*_ = 4.4206, *ρ*_*T*_ = 4.4372, *s* = 0.5, *n* = 3, *D* = 0.01, and *D*_*i*_ = 10. **(e)-(h)** Steady-state solutions to the PDE at an alternative parameter set. **(e)** The HSSs, used as initial conditions (ICs). **(f)** Starting from the Rac dominated HSS, a Rac-dominated, yet polarized pattern appears. **(g)** A balanced polarized pattern emerges from a perturbation from the coexistence HSS. **(h)** A Rho-dominated, but polarized cell results from the Rho HSS. Parameters as in (a)-(d) except *a* = 4.624, *b* = 0.7307, *c* = 0.0864, *R*_*T*_ = 1.2548 and *ρ*_*T*_ = 1.2676.

Additional PDE simulations at a different parameter set show that not only can a single parameter set give rise to different states, but that different polarized states are actually possible. Figure 3(f)-(h) illustrates that for a single parameter set, different initial conditions for the levels of active proteins (depicted by (e)) can give rise to different polarities. For example, if the system begins in a uniformly high Rac, low Rho state, it will evolve to a polarized state characterized by uniformly high but polarized Rac activity along with polarized Rho activity (e.g. panel (f)). Alternatively, if it begins in a balanced state where Rac and Rho are at similar activity levels, it will evolve to a more balanced polarity state (e.g. panel (g)). Each of these polarized states would also be linked to a different cell morphology. In the case of (f), a uniformly high Rac, but still polarized state would be associated with a large, polarized morphology. The uniformly low Rac, polarized state in (h) would be associated with a smaller, more contracted polarity state. Finally, the more balanced polarity state in (g) would correspond to an intermediate polarized state. Importantly, each of these polarity states stems from a single parameter set. Furthermore, each of the states depicted in Figure 3 maps directly onto cell morphologies that have been observed in cell populations [1, 2, 5].

In conclusion, this simple model combining known aspects of GTPase regulation gives rise to a variety of possible steady-state protein distributions, both homogeneous and heterogeneous. Furthermore, a single parameter set can give rise to a variety of possible steady-states. Thus it is possible that the wide diversity of morphologies of cells drawn from a single source may arise not from diversity of the underlying cells, but rather result from a population of similar cells exploring a rich morphological state space. That is, two cells may look different not because they have different underlying “parameters”, but because they simply ended up in different parts of the underlying state space generated by a single (or similar) set of parameters.

### 2.4. Parameter space characterization

Here we analyze the full structure of the parameter space for this GTPase regulatory system (Figure 4). We generated a sample of 10^6^ parameter sets through Latin Hypercube sampling. Well-mixed analysis is used to classify each parameter set based on the number of linearly stable homogeneous steady-states (HSSs) present as monostable, bistable, or tristable. As expected, we find that each of these three regimes are present: approximately 89% of the parameter sets are monostable, 9% bistable, and 2% tristable (Table 1). In the bistable case, the two HSSs are respectively either Rac or Rho dominated (e.g. Figures 3(a,d)). In the tristable case, both these are present along with a balanced state, namely, the coexistence HSS, where Rac and Rho are at uniform and comparable levels (Figure 3(b)). We note that the relative sparsity of multi-stable parameter sets largely results from multi-stability being restricted to lower values of the basal activation rate (*c*, see Figure 4(a)) and that a lower ceiling on that parameter could significantly effect these fractions. None the less, mono-stability is most prevalent as expected.

**TABLE 1.**
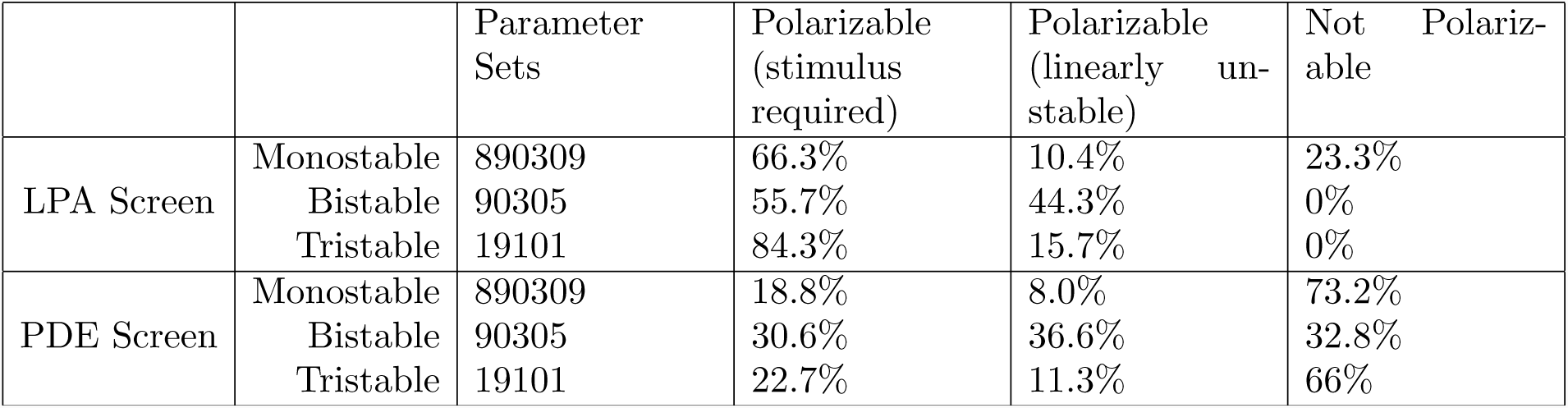
Well-mixed, LPA, and PDE Parameter screen results. From an initial 10^6^ parameter sets, the number of mono-, bi-, and tri-stable was determined (285 parameter sets were omitted due to numerical errors). “Polarizable (stimulus required)” refers to parameter sets that require a sufficiently large perturbation from a homogeneous steady state to polarize. “Polarizable (linearly unstable)” refers to those parameter sets which are LPA Unstable (LPA Screen) or those parameter sets which are unstable with respect to noise around one or more HSS in the PDE Screen. “Not Polarizable” refers to parameter sets which did not polarize using our numerical parameters and initial conditions.

**FIGURE 4.**
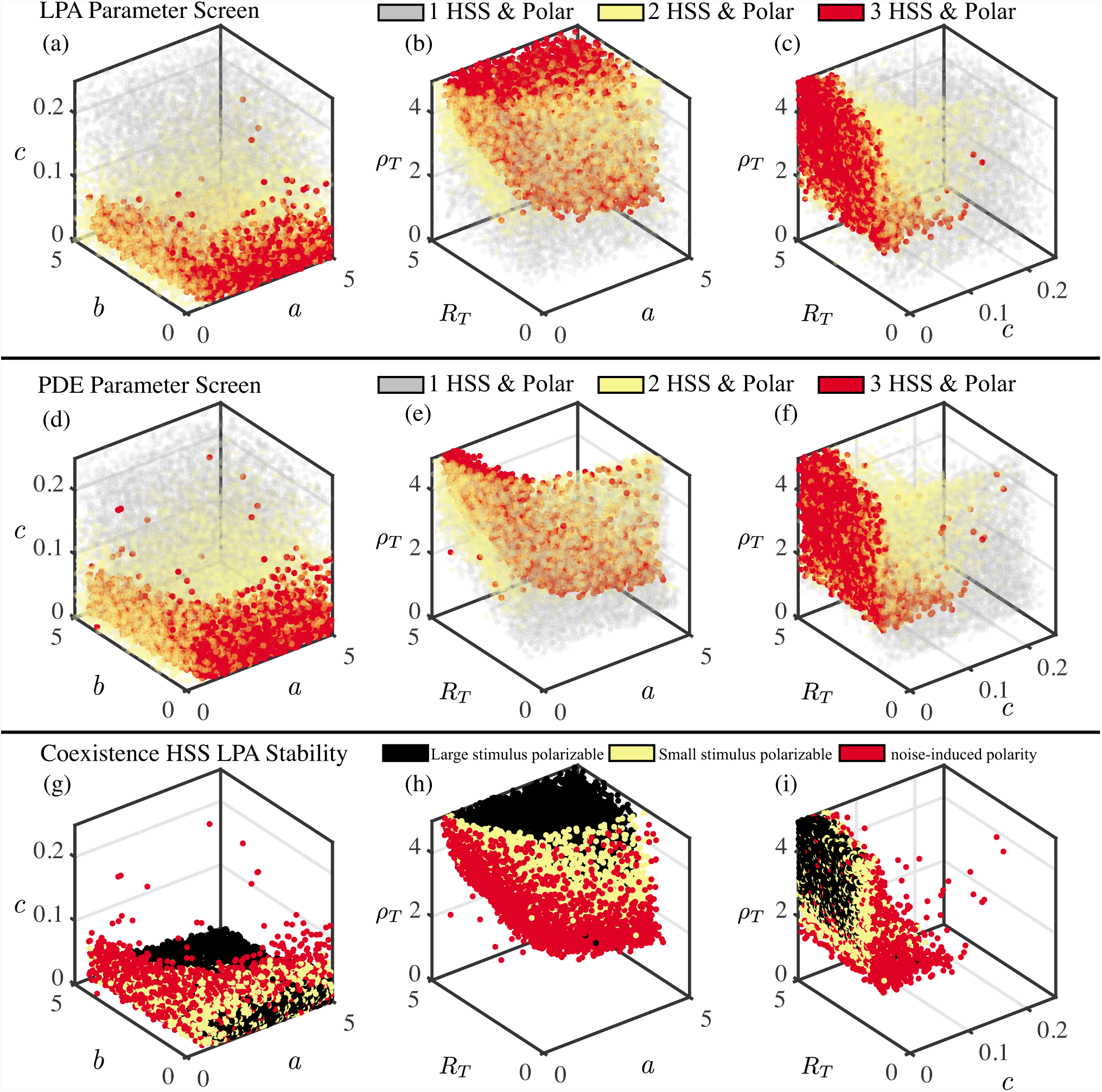
Unsupervised simulation screen reveals a nested parameter space structure. **(a)-(c)** The LPA Parameter Screen reveals structure to parameter space, with tristable polarizable (red) parameter sets nested inside bistable polarizable (yellow) parameter sets nested inside a large region of monostable polarizable (grey) parameter sets. **(d)-(e)** The PDE Parameter Screen confirms the parameter space structure predicted by the LPA Parameter Screen (plotting conventions same as (a)-(c)). **(g)-(i)** For each tristable parameter set, the level of stimulus (none, small, large) required to elicit a polarization response from the coexistence HSS is determined. For yellow points, a stimulus double (or half) the value of the HSS active protein concentration yields a response. For black points, a stimulus greater than ten times (or one tenth) the value of the HSS concentration yields a response. When the coexistence HSS is LPA unstable (red points), any perturbation away from the coexistence HSS may result in polarization. Other parameters are *s* = 0.5 and *n* = 3.

We next assessed the capacity for polarization in these mono-, bi-, and tri-stable parameter regimes. Specifically, we simulated the evolution of a spatially localized pulse-like perturbation using the LPA system of ODEs and assessed whether it was possible for that perturbation to achieve a stable “local” state (see §4 for further detail). This approach essentially allows us to easily (but approximately) ask, will the parameter set under consideration respond to a spatial stimulus, and if so, does that response require a sufficiently large stimulus or result from linear instability. This analysis indicates a large proportion of parameter sets may have the potential to elicit additional polarization states; 76.7% of monostable, 100% of bistable, and 100% of tristable parameter sets are flagged as polarizable (Table 1). Strikingly, this approach suggests that all multi-stable parameter sets *may* have the capacity for polarity. We say *may* since the results of the LPA are only approximate and require numerical PDE simulations for verification (see subsequent results). Nonetheless, these results suggest that the polarization is common in the sampled parameter ranges and actually more prevalent among multi-stable parameter sets than mono-stable parameter sets.

Visualization of the parameter space structure (using the LPA simulation screen results, (Figure 4(a)-(c)) reveals why the multi-stable parameter sets are all predicted to be polarizable. The parameter space of this model has a clear nested structure. The nucleus of this parameter space (red points indicate tristability with polarization) is a regime where a fixed parameter set can give rise to a diverse array of spatial morphologies. This is nested in a parameter regime where heterogeneity is more restricted (yellow points indicate bistability with polarization). This is in turn nested within a much more wide spread and ubiquitous parameter regime where polarity is possible but where each parameter set emits only a single HSS along with polarity (grey points indicate monostability with polarization). Thus, consistent with the quantification results in Table 1 and prior analysis [36], the multi-stable parameter regimes appear to lie within a broader polarizable regime.

To verify the predictions of the LPA, we repeated the parameter screen with full PDE numerics. Previous studies have suggested that the LPA predicts that parameter regimes are larger than they are in the actual PDEs [38]. In other words, the LPA conservatively predicts parameter regimes to be larger than they actually are. It would thus be most efficient to use the LPA to pare down the number of parameter sets for which PDE simulations are performed. However this approach has not been fully validated and thus we simulate the PDEs for all parameter sets. For each parameter set, we assessed whether or not each HSS is stable with respect to noise and whether a polarized pattern can be found. Specifically, to assess whether each HSS is stable or unstable we added noise to each HSS as an initial condition and observed the response. To determine if a particular parameter set will polarize in response to a sufficiently strong perturbation, we used a polarized pattern as an initial condition and assessed whether the system evolved to a polarized or homogeneous state. While there are differences between the PDE and LPA results, the qualitative conclusions are consistent (Figure 4(d)-(f)). First, an array of individual parameter sets can still give rise to diverse steady state spatial distributions of protein activation. Second, the nested parameter space structure is still observed. Third, polarization appears to be more common in multistable regimes than monostable regimes.

The main discrepancy between the PDE and LPA results is that there are many parameter sets where the LPA predicts polarization is possible but where PDE simulations do not achieve polarity. This is not surprising since prior studies using the LPA have generally found that while the LPA preserves the general underlying parameter space structure of a model, it generally predicts that parameter regimes are larger than they are for the PDEs. This discrepancy appears to be most prominent in the tristable and polarizable case where the PDE analysis suggests that large proportion, 65.9%, of tristable parameter sets do not give polarized patterns while the LPA Screen suggests that nearly all, 98.9%, of parameter sets are polarizable. Inspection of Figure 4(b) and (e) suggests that there is a particular region of the tristable parameter space that the LPA appears to falsely predict is polarizable. More generally, further mining of the LPA and PDE screen results shows that approximately 48% of parameter sets predicted to polarize by the PDE fail to do so when simulating the PDEs. However, effectively all parameter sets that polarize in PDE simulations are predicted to do so by the LPA; the LPA failed to predict polarization of only 0.04% of parameter sets that actually polarize in PDE simulations. Combined, these results demonstrate that the LPA is conservative in its prediction of interesting dynamics in the sense that it detects essentially all dynamically interesting parameter sets, but generally predicts regimes of behavior to be larger than they are for the PDEs.

Based on the discrepancy between the LPA and PDE results and the general interest in this parameter regime that generates a rich state space, we next focused on the regime predicted to be tristable and polarizable by the LPA. Specifically, we assessed (Figure 4(g)-(i)) whether the coexistence HSS of tristable parameter set was linearly unstable (red points) and thus would polarize in response to any stimulus, stable but easy to polarize with a relatively small stimulus (yellow points), or strongly stable requiring a large stimulus for polarization (black points). The LPA unstable points form a “shell” around the LPA stable points, suggesting that it is more difficult for the system to polarize if its parameters are further into the tristable regime. Interestingly, comparison of Figure 4(b), (e), and (h) shows that the “hard to polarize” tristable sets match up with those that were falsely predicted to polarize by the LPA. This suggests that parameter sets that reside deeper in the tristable regime are more stable against spatial perturbations, and that this point is the source of the discrepancy between the PDE and LPA results.

In conclusion, these results reveal that polarization is fairly ubiquitous throughout parameter space. The parameter space has a well defined nested structure where some regions yield more diversity of states (e.g. the number of possible steady-states for a given parameter set) than others. These results also demonstrate that the diversity of potential morphological states shown in Figure 3 is not an artifact of a single parameter set but is rather indicative of an entire regime of behavior that lies at the very core of this parameter space. Finally, these results also demonstrate the power of this approach to mapping the structure of the parameter space of this model. While there are discrepancies between the LPA (which is much more computationally efficient) and PDE (which is more accurate) approaches, the qualitative conclusions are generally consistent.

### 2.5. Numerical bifurcation analysis

To supplement our computational approach, we used numerical bifurcation analysis to study the bifurcation structure of this model. This alternative approach (1) provides another method to study parameter space structure and (2) provides a direct comparison to bifurcation analysis in previous work [36].

To perform this analysis, we first convert the GTPase model to a system of LPA ODEs. The numerical continuation package MatCont [45] is then used to perform a bifurcation analysis of that reduced system. This allows us to directly compare the dynamics of this system to that of the simpler systems consisting of only mutual-antagonism or auto-activation in [36]. Results show that the structure of the bifurcation diagram (Figure 5(a)) is qualitatively similar to that of the mutual-antagonism model (Figure 2(a)) in the sense that regimes of coexisting multi-stability and polarization are present. However, the inclusion of auto-activation can result in tristability and the formation of the coexistence HSS as in Figure 5(a). Depending on model parameters, this bifurcation diagram can take on different forms with, for example the coexistence HSS being stable or unstable (compare Figure 5(a) and 5(b)). Detailed bifurcation analysis of this system is beyond the scope of this article. This does however provide an alternative approach to demonstrate the presence of rich regimes of behavior with diverse, coexisting GTPase activation states that map onto different cell morphologies.

**FIGURE 5.**
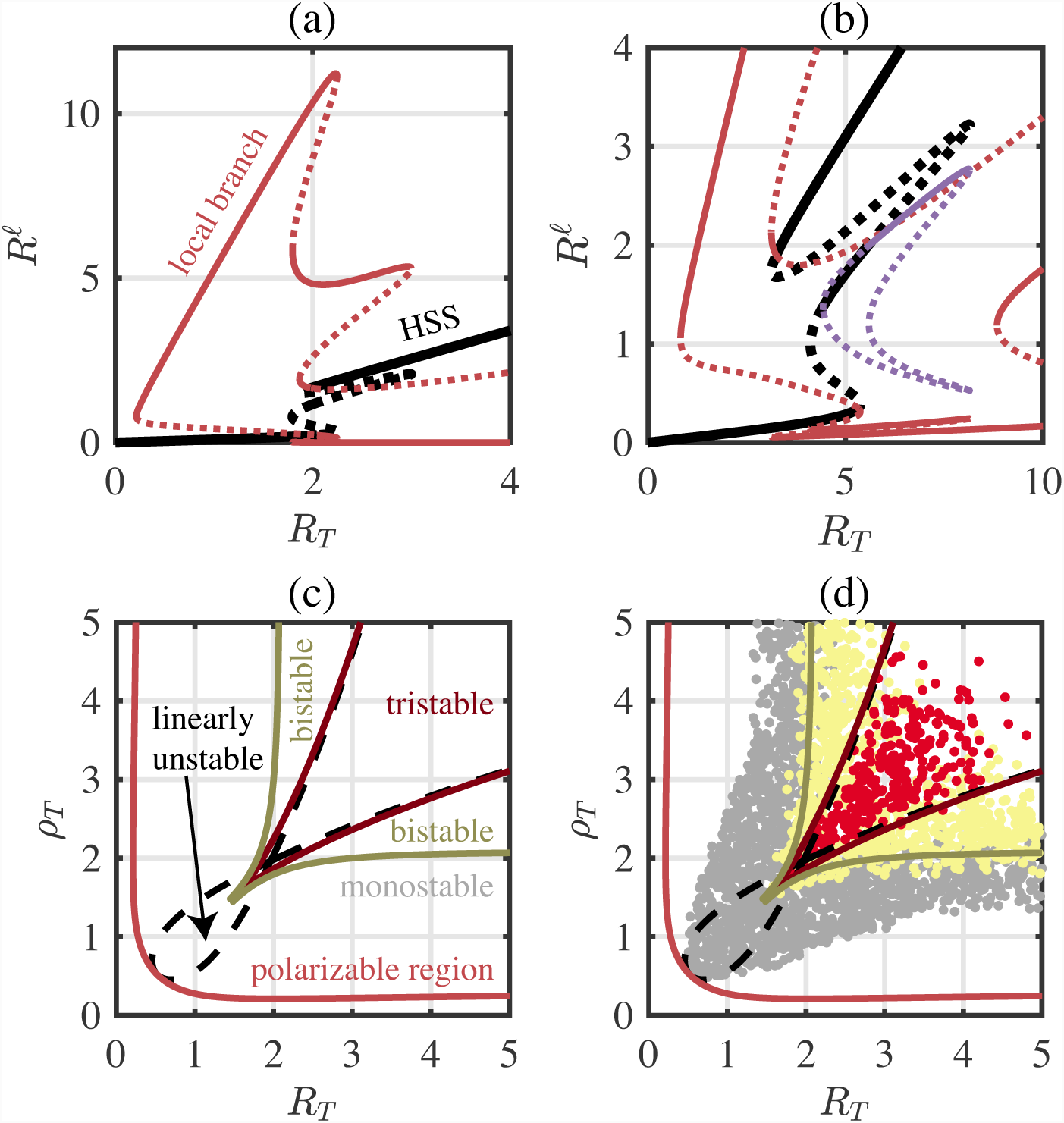
LPA bifurcation diagrams confirms parameter space structure. **(a)-(b)** The Rac global (black) and local (red and purple) solution branches are plotted with their stability (stable → solid; unstable → dashed) with respect to *R*_*T*_. **(a)** The tristable regime sits within a region of polarizability. Parameters as in Figure 2(b). **(b)** Alternative parameter space structure for different parameters that emit different stability profiles (the middle HSS is stable for some *R*_*T*_ values) and local branch structure. Parameters as in Figure 3(a)-(d). **(c)** Two-parameter bifurcation diagram of (a) with respect to *ρ*_*T*_. The region of tristability sits within a larger region of bistability, which sits within a larger region of polarizability. The red curve is a continuation of the fold bifurcation with lowest *R*_*T*_ of the local solution branch from (a). The dark red and dark yellow curves are fold continuations from the well mixed branch, colored to match the bistable and tristable points plotted in (d). The black dashed curves are the continuation of Turing bifurcations (the branch points where the local solution branches bifurcate from the global HSS solution branch). See §4 for further details of the bifurcation analysis. **(d)** A slice of the 5D parameter space from the PDE screen (Figure 4(d)-(f)) is plotted in (*R*_*T*_, *ρ*_*T*_)-plane. To match the parameters used to produce the bifurcation diagram in (a) and (c), only polarizable parameter sets with 1.3 < *a* < 2.3, 3.5 < *b* < 4.5 and 0 < *c* < 0.025 from the PDE screen are plotted. Red points: tristable and polarizable parameter sets. Yellow points: bistable and polarizable. Grey points: monostable and polarizable.

Motivated by the previously observed nested parameter space structure and by the previous bifurcation analysis in [36], we next examined the bifurcation structure in the (*R*_*T*_, *ρ*_*T*_)-plane (Figure 5(c)). We again found the nested parameter space structure with the region of polarizability encompassing both the bistable and tristable regimes. Figure 5(c) shows the results of a two-parameter continuation of the bifurcation points that delineate the transition between different regimes (i.e. two-parameter continuations of fold and branch point bifurcations, see §4 for more detail). Results once again show the tristable regime is nested within a bistable regime, which is itself nested in a broader polarizable region, confirming prior results.

Lastly, as additional verification of the parameter space structure, we projected a slice of the data from the PDE parameter screen (Figure 3(d)-(f)) onto the two-parameter bifurcation diagram (Figure 5(d)). The red points indicate tristable and polarizable parameter sets; yellow points are those parameter sets that are bistable and polarizable; and the grey points are those parameter sets that are monostable and polarizable. We find agreement between the two-parameter bifurcation diagram and the PDE parameter screen lending further credibility to our computational approach. We do note once again that while the agreement is not perfect, the regimes of behavior predicted to yield interesting dynamics by the LPA are generally larger than actual regimes found through brute-force PDE simulation. Thus, the LPA appears to capture all interesting regimes of behavior, though it does over-predict the size of those regimes.

In conclusion, numerical bifurcation results confirm the nested parameter space structure and coexistence of polarization in the monostable, bistable and tristable regimes that was found using the LPA and PDE parameter screens. Moreover, the LPA bifurcation structure in Figure 5(a) has similar qualities to that in Figure 1(a), suggesting that the addition of auto-activation to the previously studied mutual-antagonism and conservation system (as in [36]) adds to the structure of the parameter space rather than fundamentally altering it. More specifically, the addition of auto-activation leads to the genesis of a tristable and polarizable parameter regime nested within the bistable and polarizable regime that forms the nucleus of the parameter space.

## 3. DISCUSSION

Past quantification of cell shape diversity among various types of cells have revealed three essential observations. First, cells obtain a relatively restricted and discrete set of morphologies. Second, cells of different types exhibit a similar and relatively restricted set of morphologies. Third, cells of the same type can exhibit diverse cell shapes despite being derived from the same lineage. Our work here provides a hypothesis for how this landscape of discrete cell morphologies may arise and how seemingly similar cells can exhibit the full spectrum of that diversity.

In this study, we used computational modeling and analysis to demonstrate that the simple and known dynamics of Rho GTPase signaling can explain a significant amount of cell shape diversity found in cellular populations. The simple reaction-diffusion model including auto-activation, mutual-antagonism, and biochemical conservation recovers at least 6 morphologies observed in recent studies (from [2] for example). Interestingly, it is not simply the case that different regions of parameter space yield different morphologies. Rather, individual states of the model (i.e., parameter sets) can yield a variety of different shapes.

These results suggest that (1) Rho GTPase dynamics alone can explain much of the diversity of cell shapes observed experimentally and that (2) diverse morphologies may arise even in the absence of any intrinsic differences between cells. The dynamics of Rho GTPase regulation can thus explain both the presence of the discrete morphological landscape observed in a variety of cell populations as well as how populations of seemingly similar cells from the same source can explore that full landscape.

To facilitate exploration of the dynamics of this spatial model of GTPase dynamics, we further developed a new approach to efficiently and broadly explore the parameter space of this model to determine where different types of morphologies would be expected to appear. Thus, rather than pick a “representative” parameter set, we explored the entirety of the parameter space to determine the full scope of dynamics possible. Results reveal a highly structured parameter space where different parameter regimes exhibit varying levels of diversity of cell morphology. In particular, the model’s parameter space has a nested structure where the nucleus is comprised of parameter sets that can individually give rise to multiple spatial protein activation states that correspond to a variety of different shapes. Lower diversity of states is found further from the nucleus of this nested parameter space.

These results confirm conclusions of large-scale statistical analyses of cell-shape data sets that have shown the presence of cell shape “attractors” [2] and hypothesized that transitions between these attractors are may be driven by environmental perturbations that affect GTPase signaling [4]. These results however extend those prior studies. While prior studies (e.g. [2]) showed (through RNA interference [46] or chemical inhibitors [3]) that cells’ morphologies are determined by the balance between Rac and Rho in different cellular compartments, our results demonstrate for the first time that well-established cross-talk interactions between Rac and Rho can generate the full spectrum of spatial profiles necessary to generate the array of shapes found in imaging data.

The structured parameter space we observe also has similar properties to the structure found in prior mathematical analyses of spatiotemporal GTPase dynamics. In [36] and [47], bifurcation analyses illustrated a similar nested structure to parameter space. Results here however differ from those of prior studies in two important ways. First, we explore a more complete model of Rho GTPase crosstalk here that accounts for both auto-activation and mutual-antagonism rather than only mutual-antagonism. This yields a more complex and structured parameter space. Second, we have utilized unsupervised parameter space screening (similar to that found in [48, 49, 50, 51]) method based on Local Perturbation Analysis [38,39] to generate a more comprehensive understanding of the model’s dynamics and structure.

Further work is needed to understand the link between cell signaling and cell morphology. In this study, we have focused only on the core signaling dynamics of Rho GTPases to assess the types of spatial protein distributions that those dynamics generate. Simulations of spatial cell morphologies based on these distributions in 2D [52, 53, 54] or 3D [55] are needed to determine at a more detailed level how signaling characteristics are linked to morphology. Alternatively, extensions to account for the broader signaling network (e.g., [56]) regulating cytoskeletal remodeling will be required to understand how the regulation of other molecular regulators (GEFs and GAPs for example) augment cells preference for different states.

Despite these limitations, this study demonstrates that the known regulatory interactions between these proteins can (at a qualitative level) generate the full landscape of observed morphologies found in imaging studies. Furthermore detailed analysis of the model’s parameter space structure shows that these morphologies co-exist in parameter space, explaining how similar cells of the same fate drawn from a single population can exhibit a wide variety of differing morphologies. This study also validates a new highly-efficient approach for analyzing the spatial dynamics of this type of protein signaling system, that will pave the way for studying more biologically detailed models of this type in the future. These results demonstrate that Rho GTPases form the core of a cytoskeletal regulatory system governing cell shape, further supporting the picture that they act as a central signaling hub that determines how cells respond to their environmental context.

## 4. MODELS AND METHODS

Here we describe the spatial mutual-antagonism and auto-activation model of GTPase dynamics and provide more detail about our computational approach.

### 4.1. PDE model

A system of reaction-diffusion partial differential equations (PDEs) are used to model Rac and Rho GTPase activity in the cell. We track the activity of active Rac 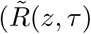 and Rho 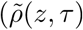, and inactive 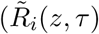 and Rho 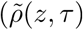. Active forms are bound to and diffuse in the cell membrane while inactive forms are free to diffuse in the cytosol. We consider the spatial domain to be a 1D slice along the cell’s diameter and assume no-flux boundary conditions. The PDEs governing the GTPase dynamics on *z* ∈ [0, *L*] and for time *τ* ∈ [0, ∞) are

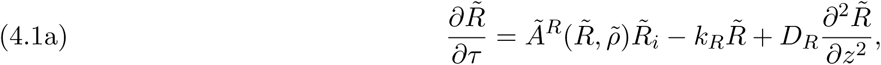

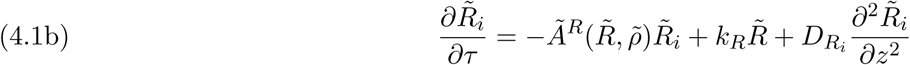

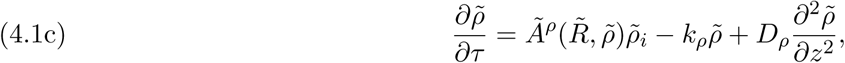

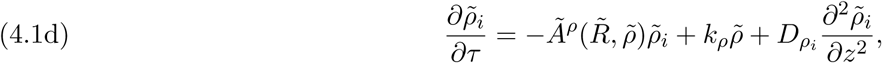

with no-ux boundary conditions

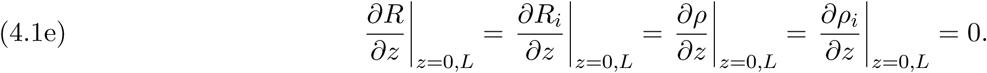

Here, *D*_*R*_ and *D*_*ρ*_ are the diffusion coefficients for active Rac and Rho, and 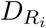 and 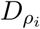 are the diffusion coefficients of the inactive forms, respectively. Since the active forms are membrane bound, the corresponding diffusion coefficients are much smaller than the inactive forms’ diffusion coefficients. We assume that the inactivation rates, *k*_*R*_ and *k*_*ρ*_, do not depend on Rac and Rho activity. Instead, we assume that mutual-antagonism and auto-activation occur through the activation rates, 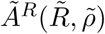 and 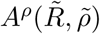:

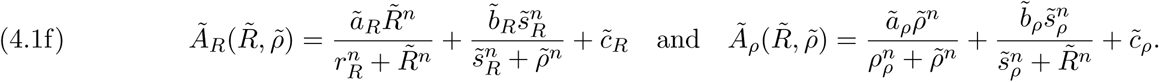

Here, the activation rates are comprised of increasing or decreasing Hill functions with exponent *n*. The parameters 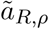 and 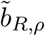 prescribe the strength of the auto-activation and mutual-antagonism, respectively, and 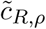 is the basal rate of activation in the absence of feedback. The parameters *r*_*R,ρ*_ and 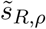 describe the amount of GTPase needed to reach the half-maximal auto-activation and antagonistic effect on the other GTPase, respectively.

Since the GTPases simply switch between active and inactive forms yet are not created nor destroyed, the activity levels of Rac and Rho must satisfy conservation statements. Indeed, adding and integrating over the domain equations (4.1a) and (4.1b), and (4.1c) and (4.1d) gives

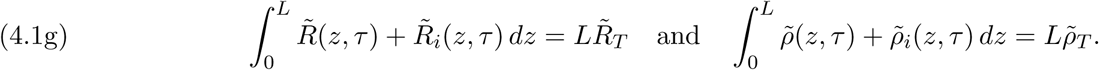

Here, 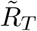 and 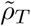 are the total Rac and Rho activity levels.

To reduce the number of parameters in the model, we scale the GTPase amounts by the corresponding half-maximal auto-activation parameter, time by the de-activation rate of Rac, and space by the cell-length:

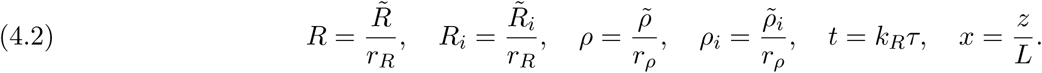

Under this scaling, the equations become

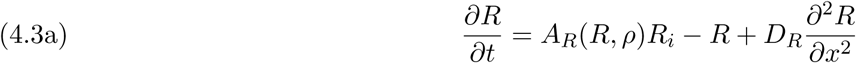

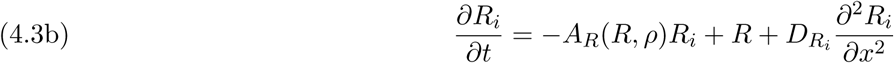

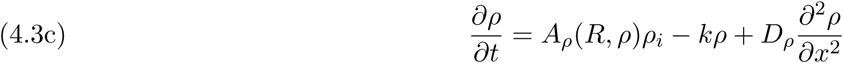

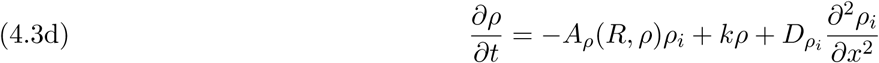

with *x* ∈ [0,1] and *t* ∈ [0, ∞). The scaled activation rates are

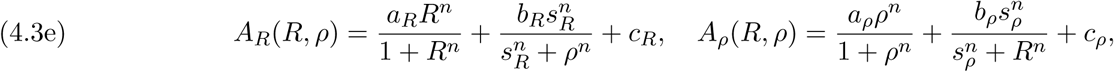

and the activity levels satisfy the conservation statements

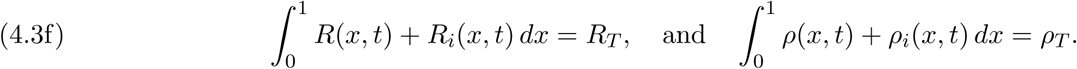

The scaled parameters are

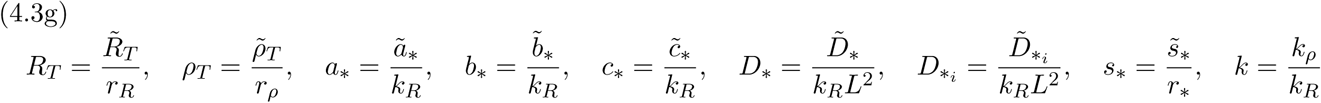

where * = *R* or *ρ*.

Finally, to reduce the complexity of the model, we restrict our attention to the case *a*_*R*_ = *a*_*ρ*_ = *a, b*_*R*_ = *b*_*ρ*_ = *b, s*_*R*_ = *s*_*ρ*_ = *s, c*_*R*_ = *c*_*ρ*_ = *c* and set *k* = 1 throughout. We also assume that the diffusion coefficients in the active state and in the inactive state are similar between Rac and Rho GTPases, so that *D*_*R*_ = *D*_*ρ*_ = *D* and that 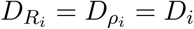.

### 4.2. Well-mixed model

In the well-mixed model, we ignore space and consider GTPase activity to be homoge-nous within the cell. This reduces the equations to a system of 4 ODEs, describing the temporal dynamics of GTPase activity.

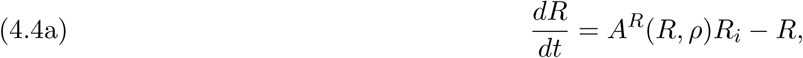

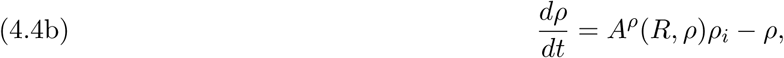

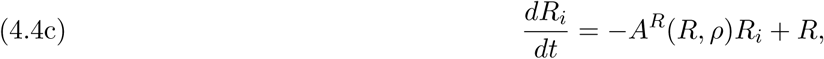

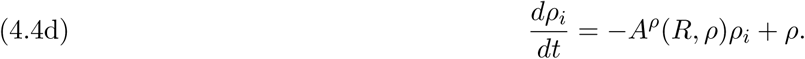

Note that this system can be reduced to a system of two ODEs which is more readily studied (for example, in the phase-plane), using the fact that the total amount of each GTPase is assumed to be constant. This means that the amount of inactive GTPase, *R*_*i*_ and *ρ*_*i*_, can be calculated as *R*_*i*_ = *R*_*T*_ − *R* and *ρ*_*i*_ = *ρ*_*T*_ − *ρ*. The reduced well-mixed model is therefore:

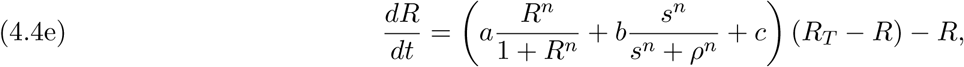

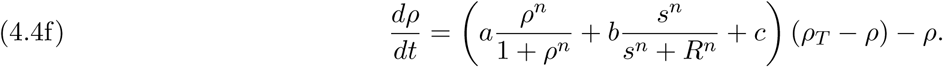

The steady-states of the well-mixed model correspond to homogeneous steady-states (HSS) of the PDE model.

### 4.3. Local Perturbation Analysis

The Local Perturbation Analysis (LPA), discussed in further detail in [37, 31, 38, 39, 44] is an approximation method that facilitates the prediction about how a spatial reaction-diffusion model will respond to spatially heterogeneous perturbations. Specifically, this method is used to assess how a homogeneous steady-state (HSS) of a PDE system comprised of fast and slow diffusing variables will respond to a spatially localized pulse-like perturbation. The fast/slow diffusion discrepancy is then exploited to describe the evolution of that pulse-like perturbation by a collection of ODEs that describe the evolution of concentrations near to (local variables) and away from (global variables) the perturbation.

Applying this reduction method and using conservation to further simplify the system yields the following equations describing the evolution of the global (*R, ρ*) and local variables (*R*^*ℓ*^, *ρ*^*ℓ*^):

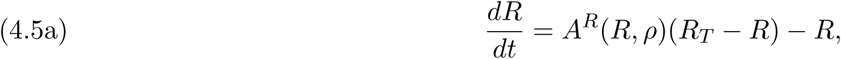

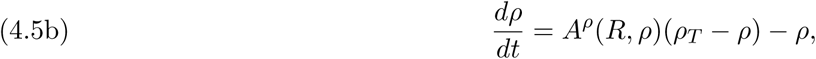

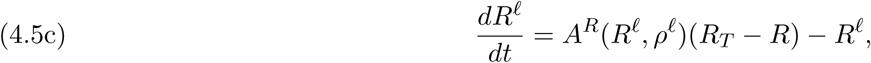

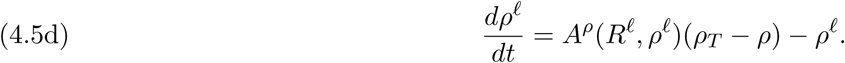

Here, the functions *A*^*R*^ and *A*^*ρ*^ capture the mutual-antagonism and auto-activation as before.

This LPA-ODE system is used in two ways. First, it is simulated to determine whether a sufficiently large perturbation will yield a response. To do this, we set the initial conditions of the global variables *R, ρ* to their HSS values (determined by simulating the well-mixed system) and then apply different initial conditions to the local variables *R*^*ℓ*^, *ρ*^*ℓ*^. If the values of *R*^*ℓ*^, *ρ*^*ℓ*^ diverge to a new value different from the HSS, this indicates a response has occurred. Otherwise, if the values of *R*^*ℓ*^, *ρ*^*ℓ*^ converge back to the HSS, the system is stable *with respect to that perturbation*. We thus systematically apply an array of perturbations to determine if the system will respond to any of them.

In addition to simulating the LPA-ODEs directly, we also apply bifurcation analysis to them in order to assess parameter space structure with respect to one or two of the parameters (Figure 2(a) and Figure 5). See subsequent sections for further detail on this approach.

### 4.4. Computational approach details

We want to understand how GTPase dynamics and cell shape depend on the parameters in the mutual-antagonism and auto-activation model. To efficiently produce a near-random sample of the 5D parameter space of interest, (*a, b, c, R*_*T*_, *ρ*_*T*_), we used MATLAB’s Latin Hypercube Sampling command with *N* = 10^6^ sample points and the default settings. The parameter ranges chosen are 0 ≤ *a, b, R*_*T*_, *ρ*_*T*_ ≤ and 0 ≤ *c* ≤ 0.25. For each parameter set, we then successively used well-mixed analysis, LPA, and PDE numerics to understand the spatial GTPase dynamics.

First, we used the well-mixed model to find the HSS of the PDE system. To determine the steady-states of the well-mixed model, we used a Latin Hypercube Sampling method to generate 50 initial conditions (*R*_0_, *ρ*_0_) in the range 0 ≤ *R*_0_ ≤ *R*_*T*_ and 0 ≤ *ρ*_0_ ≤ *ρ*_*T*_. Using these initial conditions, we then numerically solved the well-mixed model forward in time to *t* = 100 for each IC. Using the value at *t* = 100 as a new initial condition, we repeated this integration 3 times in an effort to improve the accuracy of the steady-state. Next, we classified each parameter set as monostable, bistable, or tristable based the number of unique (within some tolerance) steady-states found for each parameter set. It is possible that there are more than 3 steady-states; however, such parameter sets are rare and not considered in this study.

Second, we used LPA to determine if the parameter set could polarize. For each parameter set, we use the LPA ODEs to characterize the LPA stability of each of the HSS. If all eigenvalues of the Jacobian matrix of the LPA ODE system evaluated at a HSS all have negative real part, the HSS is called LPA Stable. If at least one of eigenvalues has positive real part, the HSS is called LPA Unstable. For those HSS which are LPA Stable, we simulated a spatially localized pulse by perturbing the initial conditions of the local variables in the LPA ODE system. That is, we chose initial conditions for the LPA ODE system 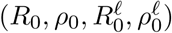 to be perturbations from the HSS. We considered large perturbations (up to 10 times the HSS amount and down to 1/10 times the HSS amount) and small perturbations (up to 2 times the HSS amount and down to 1/2 times the HSS amount) as initial conditions, and recorded if the LPA ODE system returned to the HSS or was attracted to a different steady-state. If the LPA ODE system is attracted to a different steady-state that is not another one of the other HSSs found by well-mixed analysis, for any of the HSS found for a given parameter set, we call that parameter set “polarizable (stimulus required)” (as in Figure 4(a)-(c) and Table 1). Moreover, we distinguish between those parameter sets which are polarizable with stimuli of different sizes as in Figure 4(g)-(i). Those parameter sets that are polarizable with a small stimulus, i.e., up to 2 times the HSS amount and down to 1/2 times the HSS amount, are shown as yellow points in Figure 4(g)-(i); while those parameter sets that are polarizable with a large stimulus, i.e., up to 10 times the HSS amount and down to 1/2 times the HSS amount, are shown as black points in Figure 4(g)-(i). Those parameter sets which have at least one HSS that is LPA Unstable are considered “polarizable (linearly unstable)” (Table 1) and are shown as red points in Figure 4(g)-(i).

Third, we repeated the parameter screen with PDE numerics. With PDE numerics we sought to determine: (1) whether or not a given HSS is observable (stable with respect to noise) and (2) whether a polarized pattern is a steady-state of the PDE system. To assess (1), we used each HSS as an initial condition but added noise to the active GTPase forms *R* and *ρ* (we ensured mass conservation by removing or adding the appropriate amount to the inactive forms), and checked for a polarized pattern. If one or more HSS was unstable with respect to noise, we called this parameter set “polarizable (linearly unstable)” (as in Table 1). To assess (2), we generated a polarized initial condition with *R* = *R*_*T*_ and *ρ* = 0 for 0 ≤ *x* ≤ 0.5, *R* = 0 and *ρ* = *ρ*_*T*_ for 0.5 < *x* ≤ 1, *R*_*i*_ = *R*_*T*_ */*2, and *ρ*_*i*_ = *ρ*_*T*_ */*2. If a polarized pattern results we say this parameter set is “polarizable (stimulus required)” (as in Table 1); otherwise, we say this parameter set is not polarizable. For all PDE numerics, we used MATLAB’s pdepe function with default parameters, Δ*x* = 1*/*200, Δ*t* = 1*/*400, and integrated until time *t* = 400. To check if a polarized pattern results, we computed the absolute difference between maximum and minimum for both GTPases. If this absolute difference is sufficiently large (tolerance = 0.01), we consider the pattern to be polarized.

### 4.5. Numerical bifurcation analysis

Here we describe the methods used to produce Figures 2 and 5. All numerical continuation was performed using MatCont [45] for MATLAB.

We applied standard numerical bifurcation analysis to the LPA ODEs (4.5) with *a* = 0 to produce Figure 2(a). Bifurcation analysis of the well-mixed model (4.4) was applied with *a* > 0 to produce Figure 2(b). We produced Figure 5(a) and (b) using the LPA ODEs (4.5) with parameters as in Figure 2(b) for (a) and as in Figure 3(a)-(d) for (b). Note that although Figure 5(b) has an additional local branch (shown in purple) there are common features: (1) the local branches (shown in red) have a fold (saddle-node) bifurcations for small values of *R*_*T*_; (2) the global HSS branch (shown in black) has fold bifurcations (seen clearly as the points where the blue curve changes direction and stability in Figure 2(b)); and (3) Turing bifurcations where the local branches bifurcate from the global HSS solution branch (black curves).

To gain more insight into the structure of the broader parameter space, we performed a two-parameter bifurcation analysis to track the critical bifurcations found in Figures 5(a) and (b). Specifically, we numerically continued (in the (*R*_*T*_, *ρ*_*T*_)-plane) the fold bifurcations in the local branches marking the edge of the polarizable regime, the fold bifurcations from the global branches marking the edges of the bistable and tristable regimes, as well as the Turing bifurcations marking the change in stability of the well mixed HSS in Figures 5(b). Results are shown in Figure 5(c) and copied over to Figure 5(d) without change to compare with results from the simulation screen.

## 5. ACKNOWLEDGMENTS

This work was supported by a National Science Foundation grant DMS1562078 (to WRH) and a Natural Sciences and Engineering Research Council of Canada (NSERC) Postdoctoral Fellowship Award (to CZ). This work was conducted in part using the resources of the Advanced Computing Center for Research and Education (ACCRE) at Vanderbilt University, Nashville, TN.

## REFERENCES

[1] Bakal C, Aach J, Church G, Perrimon N. Quantitative Morphological Signatures Define Local Signaling Networks Regulating Cell Morphology. Science. 2007;316(5832):1753–1756. doi:10.1126/science.1140324.

[2] Sailem H, Bousgouni V, Cooper S, Bakal C. Cross-talk between Rho and Rac GTPases drives deterministic exploration of cellular shape space and morphological heterogeneity. Open Biology. 2014;4(1):130132. doi:10.1098/rsob.130132.

[3] Byrne KM, Monsefi N, Dawson JC, Degasperi A, Bukowski-Wills JC, Volinsky N, et al. Bistability in the Rac1, PAK, and RhoA Signaling Network Drives Actin Cytoskeleton Dynamics and Cell Motility Switches. Cell Systems. 2016;2(1):38–48. doi:10.1016/j.cels.2016.01.003.

[4] Yin Z, Sadok A, Sailem H, McCarthy A, Xia X, Li F, et al. A screen for morphological complexity identifies regulators of switch-like transitions between discrete cell shapes. Nature Cell Biology. 2013;15(7):860–871. doi:10.1038/ncb2764.

[5] Cooper S, Sadok A, Bousgouni V, Bakal C. Apolar and polar transitions drive the conversion between amoeboid and mesenchymal shapes in melanoma cells. Molecular Biology of the Cell. 2015;26(22):4163–4170. doi:10.1091/mbc.e15-06-0382.

[6] Yin Z, Sailem H, Sero J, Ardy R, Wong STC, Bakal C. How cells explore shape space: A quantitative statistical perspective of cellular morphogenesis. BioEssays. 2014;36(12):1195–1203. doi:10.1002/bies.201400011.

[7] Huang B, Lu M, Jolly MK, Tsarfaty I, Onuchic J, Ben-Jacob E. The three-way switch operation of Rac1/RhoA GTPase-based circuit controlling amoeboid-hybrid-mesenchymal transition. Scientific Reports. 2014;4(1). doi:10.1038/srep06449.

[8] Nobes CD, Hall A. Rho GTPases Control Polarity, Protrusion, and Adhesion during Cell Movement. The Journal of Cell Biology. 1999;144(6):1235–1244. doi:10.1083/jcb.144.6.1235.

[9] Ridley AJ. Rho GTPase signalling in cell migration. Current Opinion in Cell Biology. 2015;36:103–112. doi:10.1016/j.ceb.2015.08.005.

[10] Wong K, Pertz O, Hahn K, Bourne H. Neutrophil polarization: Spatiotemporal dynamics of RhoA activity support a self-organizing mechanism. Proceedings of the National Academy of Sciences. 2006;103(10):3639–3644. doi:10.1073/pnas.0600092103.

[11] Xu J, Wang F, Van Keymeulen A, Herzmark P, Straight A, Kelly K, et al. Divergent Signals and Cytoskeletal Assemblies Regulate Self-Organizing Polarity in Neutrophils. Cell. 2003;114(2):201–214. doi:10.1016/s0092-8674(03)00555-5.

[12] Lin B, Holmes WR, Wang CJ, Ueno T, Harwell A, Edelstein-Keshet L, et al. Synthetic spatially graded Rac activation drives cell polarization and movement. Proceedings of the National Academy of Sciences. 2012;109(52):E3668–E3677. doi:10.1073/pnas.1210295109.

[13] Machacek M, Hodgson L, Welch C, Elliott H, Pertz O, Nalbant P, et al. Coordination of Rho GTPase activities during cell protrusion. Nature. 2009;461(7260):99–103. doi:10.1038/nature08242.

[14] Rappel WJ, Edelstein-Keshet L. Mechanisms of cell polarization. Current Opinion in Systems Biology. 2017;3:43–53. doi:10.1016/j.coisb.2017.03.005.

[15] Discher DE, Janmey P, Wang Yl. Tissue Cells Feel and Respond to the Stiffness of Their Substrate. Science. 2005;310(5751):1139– 1143. doi:10.1126/science.1116995.

[16] Vogel V, Sheetz M. Local force and geometry sensing regulate cell functions. Nature Reviews Molecular Cell Biology. 2006;7(4):265– 275. doi:10.1038/nrm1890.

[17] Houk AR, Jilkine A, Mejean CO, Boltyanskiy R, Dufresne ER, Angenent SB, et al. Membrane Tension Maintains Cell Polarity by Confining Signals to the Leading Edge during Neutrophil Migration. Cell. 2012;148(1-2):175–188. doi:10.1016/j.cell.2011.10.050.

[18] Guilluy C, Garcia-Mata R, Burridge K. Rho protein crosstalk: another social network? Trends in Cell Biology. 2011;21(12):718– 726. doi:10.1016/j.tcb.2011.08.002.

[19] Martin E, Ouellette MH, Jenna S. Rac1/RhoA antagonism defines cell-to-cell heterogeneity during epidermal morphogenesis in nematodes. The Journal of Cell Biology. 2016;215(4):483–498. doi:10.1083/jcb.201604015.

[20] Goryachev AB, Leda M. Many roads to symmetry breaking: molecular mechanisms and theoretical models of yeast cell polarity. Molecular Biology of the Cell. 2017;28(3):370–380. doi:10.1091/mbc.e16-10-0739.

[21] Ma L, Janetopoulos C, Yang L, Devreotes PN, Iglesias PA. Two Complementary, Local Excitation, Global Inhibition Mechanisms Acting in Parallel Can Explain the Chemoattractant-Induced Regulation of PI(3,4,5)P3 Response in Dictyostelium Cells. Biophysical Journal. 2004;87(6):3764–3774. doi:10.1529/biophysj.104.045484.

[22] Chau AH, Walter JM, Gerardin J, Tang C, Lim WA. Designing Synthetic Regulatory Networks Capable of Self-Organizing Cell Polarization. Cell. 2012;151(2):320–332. doi:10.1016/j.cell.2012.08.040.

[23] Park J, Holmes WR, Lee SH, Kim HN, Kim DH, Kwak MK, et al. Mechanochemical feedback underlies coexistence of qualitatively distinct cell polarity patterns within diverse cell populations. Proceedings of the National Academy of Sciences. 2017;114(28):E5750–E5759. doi:10.1073/pnas.1700054114.

[24] Holmes WR, Park J, Levchenko A, Edelstein-Keshet L. A mathematical model coupling polarity signaling to cell adhesion explains diverse cell migration patterns. PLOS Computational Biology. 2017;13(5):e1005524. doi:10.1371/journal.pcbi.1005524.

[25] Tang K, Boudreau CG, Brown CM, Khadra A. Paxillin phosphorylation at serine 273 and its effects on Rac, Rho and adhesion dynamics. PLOS Computational Biology. 2018;14(7):e1006303. doi:10.1371/journal.pcbi.1006303.

[26] Holmes WR, Liao L, Bement W, Edelstein-Keshet L. Modeling the roles of protein kinase C *β* and *?* in single-cell wound repair. Molecular Biology of the Cell. 2015;26(22):4100–4108. doi:10.1091/mbc.e15-06-0383.

[27] Holmes WR, Golding AE, Bement WM, Edelstein-Keshet L. A mathematical model of GTPase pattern formation during single-cell wound repair. Interface Focus. 2016;6(5):20160032. doi:10.1098/rsfs.2016.0032.

[28] Simon CM, Vaughan EM, Bement WM, Edelstein-Keshet L. Pattern formation of Rho GTPases in single cell wound healing. Molecular Biology of the Cell. 2013;24(3):421–432. doi:10.1091/mbc.e12-08-0634.

[29] Iglesias PA, Devreotes PN. Navigating through models of chemotaxis. Current Opinion in Cell Biology. 2008;20(1):35–40. doi:10.1016/j.ceb.2007.11.011.

[30] Verkhovsky AB. Cell Polarization: Mechanical Switch for a Chemical Reaction. Current Biology. 2012;22(2):R58–R61. doi:10.1016/j.cub.2011.12.012.

[31] Edelstein-Keshet L, Holmes WR, Zajac M, Dutot M. From simple to detailed models for cell polarization. Philosophical Transactions of the Royal Society B: Biological Sciences. 2013;368(1629):20130003. doi:10.1098/rstb.2013.0003.

[32] Holmes WR, Edelstein-Keshet L. A Comparison of Computational Models for Eukaryotic Cell Shape and Motility. PLOS Computational Biology. 2012;8(12):e1002793. doi:10.1371/journal.pcbi.1002793.

[33] Mori Y, Jilkine A, Edelstein-Keshet L. Wave-Pinning and Cell Polarity from a Bistable Reaction-Diffusion System. Biophysical Journal. 2008;94(9):3684–3697. doi:10.1529/biophysj.107.120824.

[34] Mori Y, Jilkine A, Edelstein-Keshet L. Asymptotic and Bifurcation Analysis of Wave-Pinning in a Reaction-Diffusion Model for Cell Polarization. SIAM Journal on Applied Mathematics. 2011;71(4):1401–1427. doi:10.1137/10079118x.

[35] Jilkine A, Maree AFM, Edelstein-Keshet L. Mathematical Model for Spatial Segregation of the Rho-Family GTPases Based on Inhibitory Crosstalk. Bulletin of Mathematical Biology. 2007;69(6):1943–1978. doi:10.1007/s11538-007-9200-6.

[36] Holmes WR, Edelstein-Keshet L. Analysis of a minimal Rho-GTPase circuit regulating cell shape. Physical Biology. 2016;13(4):046001. doi:10.1088/1478-3975/13/4/046001.

[37] Grieneisen V. Dynamics of Auxin Patterning in Plant Morphogenesis. University of Utrecht; 2009.

[38] Holmes WR. An Efficient, Nonlinear Stability Analysis for Detecting Pattern Formation in Reaction Diffusion Systems. Bulletin of Mathematical Biology. 2014;76(1):157–183. doi:10.1007/s11538-013-9914-6.

[39] Holmes WR, Mata MA, Edelstein-Keshet L. Local Perturbation Analysis: A Computational Tool for Biophysical Reaction-Diffusion Models. Biophysical Journal. 2015;108(2):230–236. doi:10.1016/j.bpj.2014.11.3457.

[40] Guantes R, Poyatos JF. Multistable Decision Switches for Flexible Control of Epigenetic Differentiation. PLOS Computational Biology. 2008;4(11):e1000235. doi:10.1371/journal.pcbi.1000235.

[41] Wu F, Su RQ, Lai YC, Wang X. Engineering of a synthetic quadrastable gene network to approach Waddington landscape and cell fate determination. eLife. 2017;6:e23702. doi:10.7554/elife.23702.

[42] Holmes WR, Reyes de Mochel NS, Wang Q, Du H, Peng T, Chiang M, et al. Gene Expression Noise Enhances Robust Organization of the Early Mammalian Blastocyst. PLOS Computational Biology. 2017;13(1):e1005320. doi:10.1371/journal.pcbi.1005320.

[43] Otsuji M, Ishihara S, Co C, Kaibuchi K, Mochizuki A, Kuroda S. A Mass Conserved Reaction–Diffusion System Captures Properties of Cell Polarity. PLOS Computational Biology. 2007;3(6):e108. doi:10.1371/journal.pcbi.0030108.

[44] Mata MA, Dutot M, Edelstein-Keshet L, Holmes WR. A model for intracellular actin waves explored by nonlinear local perturbation analysis. Journal of Theoretical Biology. 2013;334:149–161. doi:10.1016/j.jtbi.2013.06.020.

[45] Dhooge A, Govaerts W, Kuznetsov YA, Meijer HGE, Sautois B. New features of the software MatCont for bifurcation analysis of dynamical systems. Mathematical and Computer Modelling of Dynamical Systems. 2008;14(2):147–175. doi:10.1080/13873950701742754.

[46] Yamazaki D, Itoh T, Miki H, Takenawa T. srGAP1 regulates lamellipodial dynamics and cell migratory behavior by modulating Rac1 activity. Molecular Biology of the Cell. 2013;24(21):3393–3405. doi:10.1091/mbc.e13-04-0178.

[47] Trong PK, Nicola EM, Goehring NW, Kumar KV, Grill SW. Parameter-space topology of models for cell polarity. New Journal of Physics. 2014;16(6):065009. doi:10.1088/1367-2630/16/6/065009.

[48] Wang Y, Ku CJ, Zhang ER, Artyukhin AB, Weiner OD, Wu LF, et al. Identifying Network Motifs that Buffer Front-to-Back Signaling in Polarized Neutrophils. Cell Reports. 2013;3(5):1607–1616. doi:10.1016/j.celrep.2013.04.009.

[49] Nguyen LK, Degasperi A, Cotter P, Kholodenko BN. DYVIPAC: an integrated analysis and visualisation framework to probe multi-dimensional biological networks. Scientific Reports. 2015;5(1):12569. doi:10.1038/srep12569.

[50] Huang B, Lu M, Jia D, Ben-Jacob E, Levine H, Onuchic JN. Interrogating the topological robustness of gene regulatory circuits by randomization. PLOS Computational Biology. 2017;13(3):e1005456. doi:10.1371/journal.pcbi.1005456.

[51] Yue H, Camley BA, Rappel WJ. Minimal Network Topologies for Signal Processing during Collective Cell Chemotaxis. Biophysical Journal. 2018;114(12):2986–2999. doi:10.1016/j.bpj.2018.04.020.

[52] Vanderlei B, Feng JJ, Edelstein-Keshet L. A Computational Model of Cell Polarization and Motility Coupling Mechanics and Biochemistry. Multiscale Modeling & Simulation. 2011;9(4):1420–1443. doi:10.1137/100815335.

[53] Alonso S, Stange M, Beta C. Modeling random crawling, membrane deformation and intracellular polarity of motile amoeboid cells. PLOS ONE. 2018;13(8):e0201977. doi:10.1371/journal.pone.0201977.

[54] Camley BA, Zhao Y, Li B, Levine H, Rappel WJ. Crawling and turning in a minimal reaction-diffusion cell motility model: Coupling cell shape and biochemistry. Physical Review E. 2017; 95(1):012401. doi:10.1103/physreve.95.012401.

[55] Cusseddu D, Edelstein-Keshet L, Mackenzie JA, Portet S, Madzvamuse A. A coupled bulk-surface model for cell polarisation. Journal of Theoretical Biology. 2018;doi:10.1016/j.jtbi.2018.09.008.

[56] Kim TH, Monsefi N, Song JH, von Kriegsheim A, Vandamme D, Pertz O, et al. Network-based identification of feedback modules that control RhoA activity and cell migration. Journal of Molecular Cell Biology. 2015;7(3):242–252. doi:10.1093/jmcb/mjv017.

